# Muscular dystrophy-associated lamin variants disrupt cellular organization through a nucleolar-ribosomal axis controlling cytoplasmic macromolecular crowding

**DOI:** 10.1101/2025.09.22.677926

**Authors:** Xiangyi Ding, Sweta Kumari, Ellen Gregory, Daniel A. Starr, G. W. Gant Luxton

**Author notes:** Corresponding authors. Email: DAS or GWGL.

## Abstract

Emery-Dreifuss muscular dystrophy (EDMD) arises from mutations in nuclear lamins or emerin. Current pathological models emphasize defects in nuclear mechanics and transcription regulation. Yet these models do not fully explain the complexity of EDMD or other laminopathies. Here, we uncover an emerging pathway linking nuclear lamina defects to the reorganization of cytoplasmic biophysics, revealing how nuclear dysfunction cascades throughout the cell. Using *Caenorhabditis elegans* EDMD models, we demonstrate that lamin mutations dramatically alter cytoplasmic organization, reducing macromolecular crowding and increasing diffusivity of 40 nm <u>G</u>enetically <u>E</u>ncoded <u>M</u>ultimeric (GEM) nanoparticles. These striking biophysical changes coincide with nuclear positioning defects and collapsed endoplasmic reticulum architecture, mirroring phenotypes associated with ribosome depletion. We propose a mechanism where mutations in the *C. elegans* lamin *lmn-1* disrupt nucleolar density and ribosome biogenesis, creating a nucleolar-ribosomal axis that propagates defects from the nucleus to the cytoplasm. Genetic interactions between *lmn-1* and ribosomes support this regulatory relationship. While individual depletion of other nuclear envelope proteins produces minimal effects, combined loss of the functionally redundant emerin ortholog *emr-1* and LEM-domain protein *lem-2* phenocopied *lmn-1* mutants, demonstrating that cytoplasmic biophysical disruption lies at EDMD’s pathogenic core. Our findings establish a paradigm where nuclear lamina defects fundamentally rewire cellular biophysics through nucleolar-ribosomal dysfunction, opening transformative therapeutic avenues for treating laminopathies.

## INTRODUCTION

Emery-Dreifuss muscular dystrophy (EDMD) is a devastating neuromuscular disorder characterized by progressive skeletal muscle weakness, cardiac conduction defects, and joint contractures^1^. EDMD arises primarily from mutations in either *LMNA*, encoding nuclear A-type lamins (lamins A/C), or *EMD*, encoding the inner nuclear membrane protein emerin^1^. A type lamins confer mechanical stability to the nucleus, connect the nucleus to the cytoskeleton via linker of nucleoskeleton and cytoskeleton (LINC) complexes, and regulate chromatin organization to control gene expression^2^. Emerin, a LEM-domain protein, interacts structurally with A-type lamins and contributes to lamina functions^2^. Together, these observations highlight that EDMD arises from disruption of the lamina network rather than a single protein, underscoring the challenge of understanding how lamina dysfunction mechanistically leads to the tissue-specific EDMD pathologies.

A defining cellular hallmark of EDMD is aberrant nuclear positioning in skeletal muscle fibers^3,4^. In healthy muscle, nuclei distribute peripherally near the sarcolemma, but EDMD patients show nuclei inappropriately clustered at the fiber center^3^. This nuclear mispositioning indicates disrupted nuclear-cytoskeletal coupling, likely through LINC complexes^5,6^. However, nuclear positioning defects alone cannot fully explain EDMD’s complex pathophysiology, as other diseases with prominent nuclear mispositioning present distinct progression patterns^7^, indicating that additional cellular mechanisms must be disrupted.

Current laminopathy models focus primarily on altered mechanotransduction and dysregulated gene expression^8^. While these mechanisms contribute to disease, neither fully accounts for the broad cellular disorganization observed in EDMD tissues, including ruffled nuclear envelope and mispositioned muscle nuclei^8^. Recent advances in understanding cytoplasmic biophysical properties suggest a third possibility: lamin mutations might disrupt fundamental physical properties of the cellular interior, leading to widespread organizational defects^9,10^.

The eukaryotic cytoplasm is a highly organized, crowded environment where macromolecular interactions influence all cellular functions^11^. Macromolecular crowding affects protein folding, enzyme kinetics, liquid-liquid phase separation, and organelle positioning^11^. Ribosome concentration is a major determinant of cytoplasmic macromolecular crowding and cellular organization in yeast and mammalian cells^9^. Ribosome depletion reduces cytoplasmic macromolecular crowding, disrupts nuclear positioning, and causes endoplasmic reticulum (ER) network collapse in *C. elegans*^10^. Ribosomes assemble in the nucleolus before cytoplasmic export, creating a direct link between nuclear function and cytoplasmic organization^9,12^.

*C. elegans* provides unique advantages for investigating potential interactions between nuclear lamins, nucleolar function, and cytoplasmic macromolecular crowding. The nematode possesses a single lamin gene, *lmn-1*, whose product functionally substitutes for both A- and B-type mammalian lamins^5,13,14^. Like human lamins, LMN-1 localizes to the nuclear periphery, interacts with chromatin, and is required for proper nuclear shape and positioning^5,14,15^. *C. elegans* EDMD models engineered using CRISPR/Cas9 faithfully recapitulate many disease phenotypes, including reduced fitness, movement disorders, and nuclear morphology defects^15^. The *lmn-1(R64P)* mutation, corresponding to the severe human variant *LMNA* p.R50P, is particularly disruptive, causing both polymerization defects *in vitro*, and EDMD-like phenotypes *in vivo*^13,15^. *C. elegans* is also well suited for measuring macromolecular crowding through *in vivo* nanorheology using <u>G</u>enetically <u>E</u>ncoded <u>M</u>ultimeric nanoparticles (GEMs) ^9,10^, fluorescent protein assemblies that report local crowding conditions through their diffusive behavior in live animals. This approach revealed that cytoplasmic mesoscale macromolecular crowding in *C. elegans* tissues is regulated by the giant ER/outer nuclear membrane Klarsicht/ANC-1/SYNE homology (KASH) protein ANC-1 and ribosome concentration^10^, both of which independently influence nuclear positioning and ER morphology.

Here, we test whether EDMD-associated lamin mutations disrupt cellular organization through effects on mesoscale macromolecular crowding, ribosome biogenesis, and nucleolar function. Our findings reveal a nucleolar-ribosomal axis through which nuclear lamina defects propagate to cause broad cellular dysfunction, providing a mechanistic basis for the cellular disorganization observed in EDMD tissues.

## RESULTS

### EDMD-associated lamin variants disrupt cytoplasmic biophysical properties *in vivo*

To investigate how lamin mutations affect cytoplasmic biophysical properties in living tissues, we employed *in vivo* nanorheology in living *C. elegans*. This approach provides real-time quantification of cytoplasmic biophysical properties in intact animals using 40 nm GEMs as fluorescent tracers to explore the mesoscale intracellular environment^9,10^. We quantified cytoplasmic biophysical properties by analyzing mean squared displacement curves to calculate effective particle diffusion coefficients (*D_eff_)* through tracking trajectories of thousands of individual GEMs with high-speed spinning-disc confocal microscopy. Measurements were made in the hypodermis and intestine of homozygous *lmn-1* mutant animals and compared to heterozygous controls maintained by *hT2* balancer (**Movie S1-S2**)^15,16^. In heterozygous control animals, most GEMs were highly constrained within the crowded cytoplasm, while a smaller population of GEMs showed faster, more diffusive movements (**Figs. 1A and 1C**). This bimodal pattern was consistent with previous observations in wild-type animals and reflects the spatially heterogeneous nature of the intracellular environment^10^.

**Figure 1.**
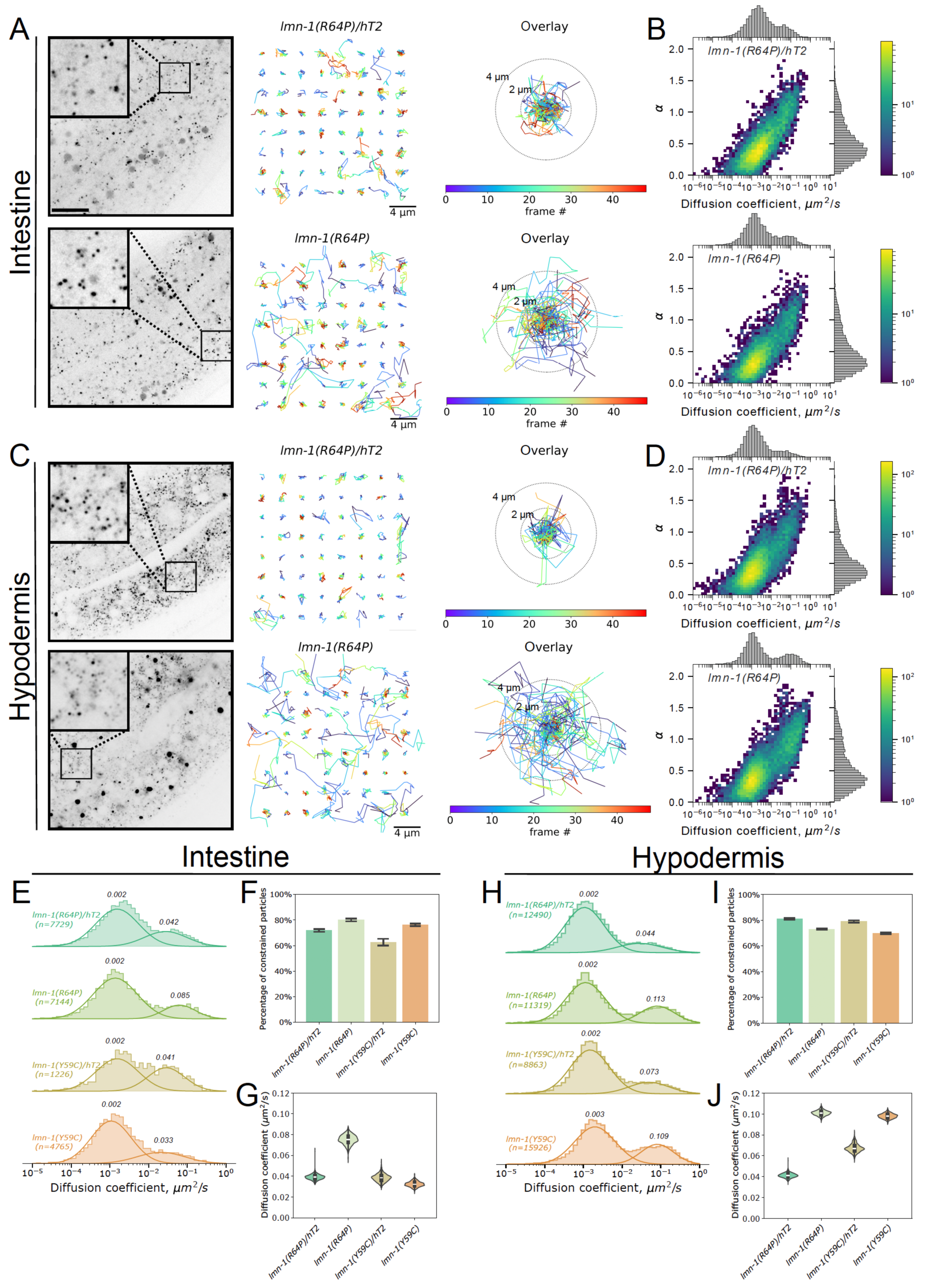
*lmn-1* variants Y59C and R64P disrupt cytoplasmic biophysical properties. **A, C)** Representative inverted greyscale spinning disc confocal images of GEMs in the intestine (A) or hypodermis (C) of day 1 *lmn-1(R64P)/hT2* (top) or *lmn-1(R64P)* (bottom) adults. Scale bar: 10 µm. Magnified insets show detailed views of boxed regions. Right panels display representative particle trajectories color-coded by frame number. Scale bar: 4 µm. Concentric circles indicate 2 μm and 4 μm radii from the trajectory origin. **B, D)** 2D probability density histograms correlating D_eff_ with the anomalous diffusion parameter (α) over the initial 100 ms for GEM diffusion in the intestine (B) or hypodermis (D) of day 1 *lmn-1(R64P)/hT2* (top) or *lmn-1(R64P)*(bottom) adults. Marginal distributions shown as frequency histograms on top and right axes. Color scale indicates particle count per bin. **E, H)** Gaussian Mixture Model (GMM) analysis of GEM *D_eff_* in the intestine (E) or hypodermis (H) comparing the indicated strains. **F, I, G, J)** Bootstrap analysis of GMM parameters comparing GEM diffusion in the intestine (F, G) or hypodermis (I, J) of the indicated strains. The percentage of constrained GEMs (F,I) with mean ± SD and the *D_eff_* values of unconstrained GEMs (G,J) are shown.

We next characterized GEM behavior in EDMD-associated *lmn-1* homozygous mutants, focusing on two early-onset variants: *lmn-1(R64P)* and *lmn-1(Y59C),* respectively corresponding to human *LMNA* p.R50P and p.Y45C^13,15,17^. The *lmn-1(R64P)* mutation causes polymerization defects *in vitro*^13^. Homozygous *lmn-1(R64P)* mutants showed substantial and reproducible increases in GEM mobility compared to balanced heterozygous controls in both the intestine and the hypodermis. In contrast, homozygous *lmn-1(Y59C)* mutants displayed marked increases in GEM mobility in the hypodermis but minimal effects in the intestine (**Figs. 1A-D, S1**). To quantitatively analyze these distinct kinetic populations, we performed mean squared displacement (MSD) analysis (**Fig. S2**) and applied Gaussian mixture modeling to objectively separate GEM trajectories into “constrained” (low mobility) and “unconstrained” (high mobility) classes, using our previously developed analytical framework ^10^ (**Figs. 1E and 1H**). Enhanced diffusion occurred specifically in the faster-moving, unconstrained particle population, while the constrained population remained largely unaffected in both fraction size and diffusion characteristics. Bootstrap analysis with 1,000 iterations revealed that *lmn-1(R64P)* markedly increased the *D_eff_* of unconstrained particles in both intestinal (**Fig. 1G**) and hypodermal tissues (**Fig. 1J**), whereas *lmn-1(Y59C)* caused pronounced increases only in the hypodermis (**Fig. 1G**). The fraction of constrained particles remained statistically similar to controls in all cases (**Figs. 1F and 1I**). These findings indicate that *lmn-1(R64P)* broadly disrupts cytoplasmic macromolecular crowding, while *lmn-1(Y59C)* exerts tissue-specific effects, particularly in the hypodermis (**Figs. 1G and 1J**). Together, these results demonstrate that EDMD-associated lamin mutations alter cytoplasmic biophysical properties by reducing mesoscale macromolecular crowding, albeit with distinct tissue sensitivities.

### EDMD-associated lamin variants recapitulate disease-relevant cellular defects

Disrupted nuclear positioning is consistently observed in the skeletal muscle of EDMD patients^1,3^. We therefore examined whether our *C. elegans* lamin mutants recapitulate this phenotype by analyzing nuclear positioning in the hypodermis, a single-cell syncytial layer containing 139 nuclei that are evenly spaced apart and anchored in place by mechanisms involving the giant KASH protein ANC-1^18,19^. We quantified nuclear anchorage defects by measuring the percentage of nuclei found in abnormal clusters^20,21^.

EDMD-associated lamin mutations caused significant nuclear clustering compared to controls (**Figs. 2A-B**). The severe *lmn-1(R64P)* variant produced the strongest effect, with ∼35% of nuclei clustered, versus <5% in wild-type animals. *lmn-1(Y59C)* had an intermediate effect, consistent with its less severe impact on cytoplasmic crowding in the hypodermis. This nuclear mispositioning defect parallels those observed in EDMD patient tissues. The severity hierarchy (R64P > Y59C) also matches known *in vitro* polymerization defects and organismal phenotypes^13,15^, demonstrating that EDMD-associated lamin mutations disrupt the mechanisms governing nuclear positioning.

**Figure 2.**
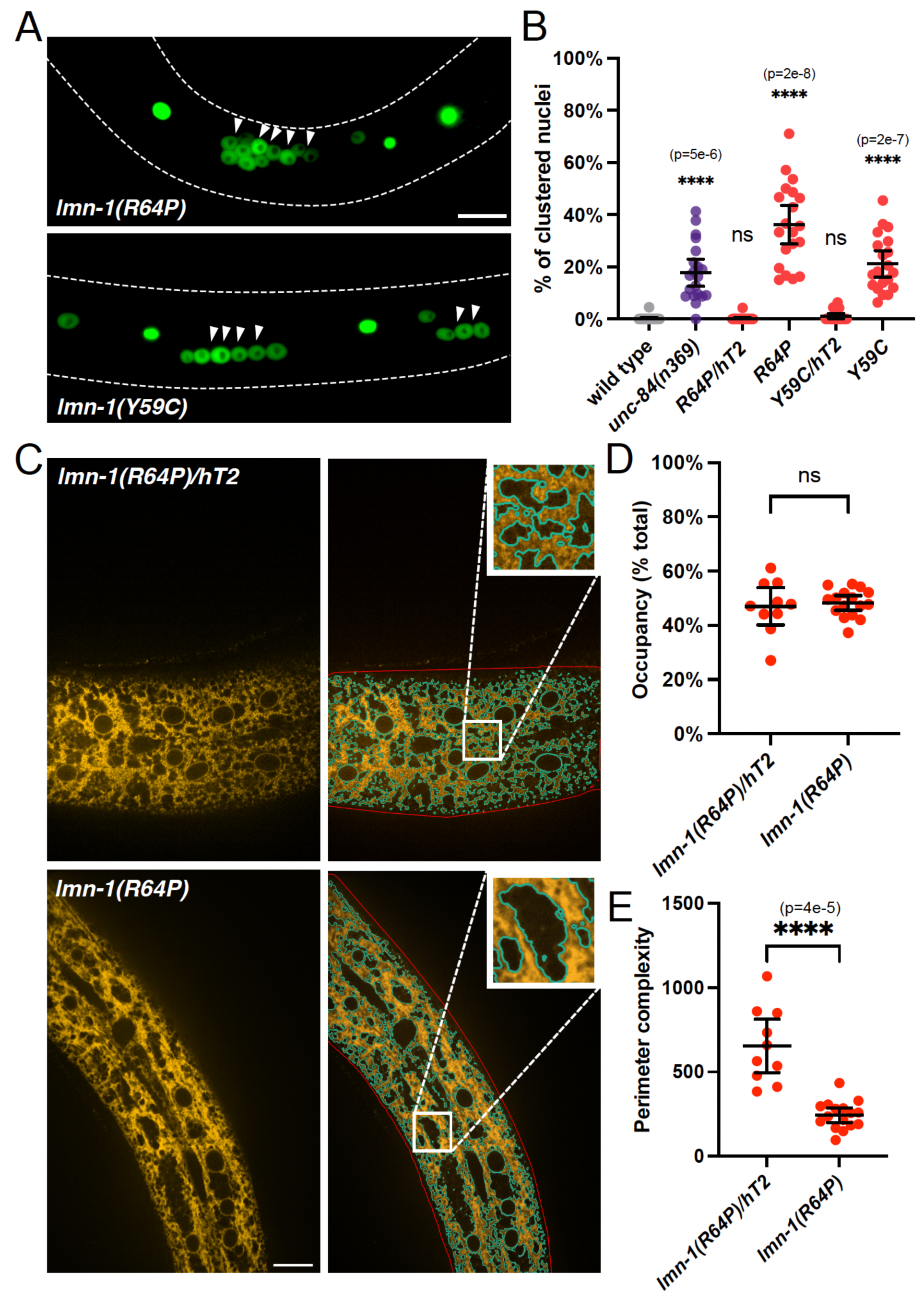
*lmn-1* variants disrupt nuclear positioning and ER morphology in the hypodermis. **A)** Representative spinning disc confocal images of GFP-labeled hypodermal nuclei (*ycIs10*, see Methods) in day 1 adult worms of the indicated strains. Arrowheads indicate defective nuclear anchoring; dotted lines outline worm boundaries. Scale bar: 10 μm. **B)** Nuclear positioning defects quantified as percentage of touching nuclei per worm in day 1 adults of the indicated strains. Each point represents one worm (n=20 for each genotype). Welch’s ANOVA test: W(5, 50) = 44, p < 0.001. **C)** Representative spinning disc confocal images of hypodermal ER organization in the indicated strains at day 1 of adulthood. Left panels: ER morphology visualized by mKate2::TRAM-1 fluorescence. Right panels: Thresholded images showing ER outline (cyan) and hypodermal boundary (red). Scale bar: 10 μm. Zoom-in insets show magnified views of boxed regions. Each point represents each worm’s ER morphological analysis. **D-E**) ER network analysis showing ER occupancy percentage (D) (n=11, 10, 16 from left to right) and perimeter complexity (see Methods) (E) (n=11, 10, 16 from left to right) across genotypes. Kruskal-Wallis test: H(2) = 19.91, p < 0.001. Statistical significance is indicated by asterisks (****: p<0.0001, ***: p≤0.001,**: p≤0.01, *: p≤0.05, ns: p>0.05) with exact p-values shown above each comparison.

To examine whether lamin mutations affect broader cellular architecture beyond nuclear positioning, we examined ER morphology. The ER forms extensive networks throughout hypodermal syncytia that depend on *anc-1* pathways involved in cytoplasmic constraint and macromolecular crowding pathways reliant on ribosome concentration^10,18^. In wild type animals, the hypodermal ER displays a complex branched network of interconnected tubules and sheets with small space between ER-rich regions (**Fig. 2C**). *lmn-1(R64P)* mutants, however, showed altered ER architecture, with larger void spaces suggesting network collapse (**Fig. 2C**). The mutant ER formed more sheet-like structures with reduced tubular branching, resembling simplified networks observed under cytoplasmic stress or crowding defects^22^. We quantified these defects using described morphometric approaches^10,23^, measuring both ER occupancy (total area coverage) and network complexity (perimeter-to-area ratio). While *lmn-1(R64P)* mutants showed no significant change in total ER area, they exhibited marked reduction in network complexity (**Figs. 2D and 2E**). Notably, the nature of ER network defects in *lmn-1(R64P)* mutants closely matched those previously observed in ribosome-depleted animals^10^, suggesting a shared underlying mechanism. These data demonstrate that lamin dysfunction produces cellular defects quantitatively and qualitatively similar to ribosome depletion effects, pointing toward a common pathway linking nuclear lamina integrity to cytoplasmic organization.

### LMN-1 and ribosomes function in a common pathway controlling cellular organization

Since *lmn-1(R64P)* mutants phenocopy several cellular organization defects previously observed in ribosome-depleted animals, including reduced cytoplasmic crowding, nuclear mispositioning, and ER network collapse, we hypothesized that lamins and ribosomes function in the same pathway to regulate mesoscale macromolecular crowding and cellular organization. This model predicts that if lamins act upstream of ribosomes in the same pathway, combining *lmn-1* mutations with partial ribosome depletion should produce epistatic rather than additive effects. To test this prediction, we performed genetic interaction studies using *rps-18(RNAi)* to partially deplete ribosomes in various *lmn-1* mutant backgrounds and quantified three independent phenotypes: GEM diffusion, nuclear positioning, and ER architecture.

Ribosome depletion by *rps-18(RNAi)* robustly increased intestinal GEM diffusion in heterozygous *lmn-1(R64P)/hT2, lmn-1(Y59C)/hT2,* and homozygous *lmn-1(Y59C)* animals, reproducing previous findings that ribosomes regulate cytoplasmic crowding (**Figs. 3A, 3C**)^10^. This response demonstrated that ribosome depletion retained its ability to disrupt cytoplasmic organization in genetic backgrounds with partial lamin function. However, *rps-18(RNAi)* failed to further increase cytoplasmic GEM diffusion in homozygous *lmn-1(R64P)* mutants, revealing a clear epistatic interaction (Figs. 1G and 3E-G). This epistasis suggests that the severe *lmn-1(R64P)* mutation maximally disrupts the lamin-ribosome pathway. The differential responses between *lmn-1(Y59C)* (additive effects) and *lmn-1(R64P)* (epistatic effects) provide insight into the severity hierarchy of EDMD variants and their relative impacts on the lamin-ribosome regulatory axis.

**Figure 3.**
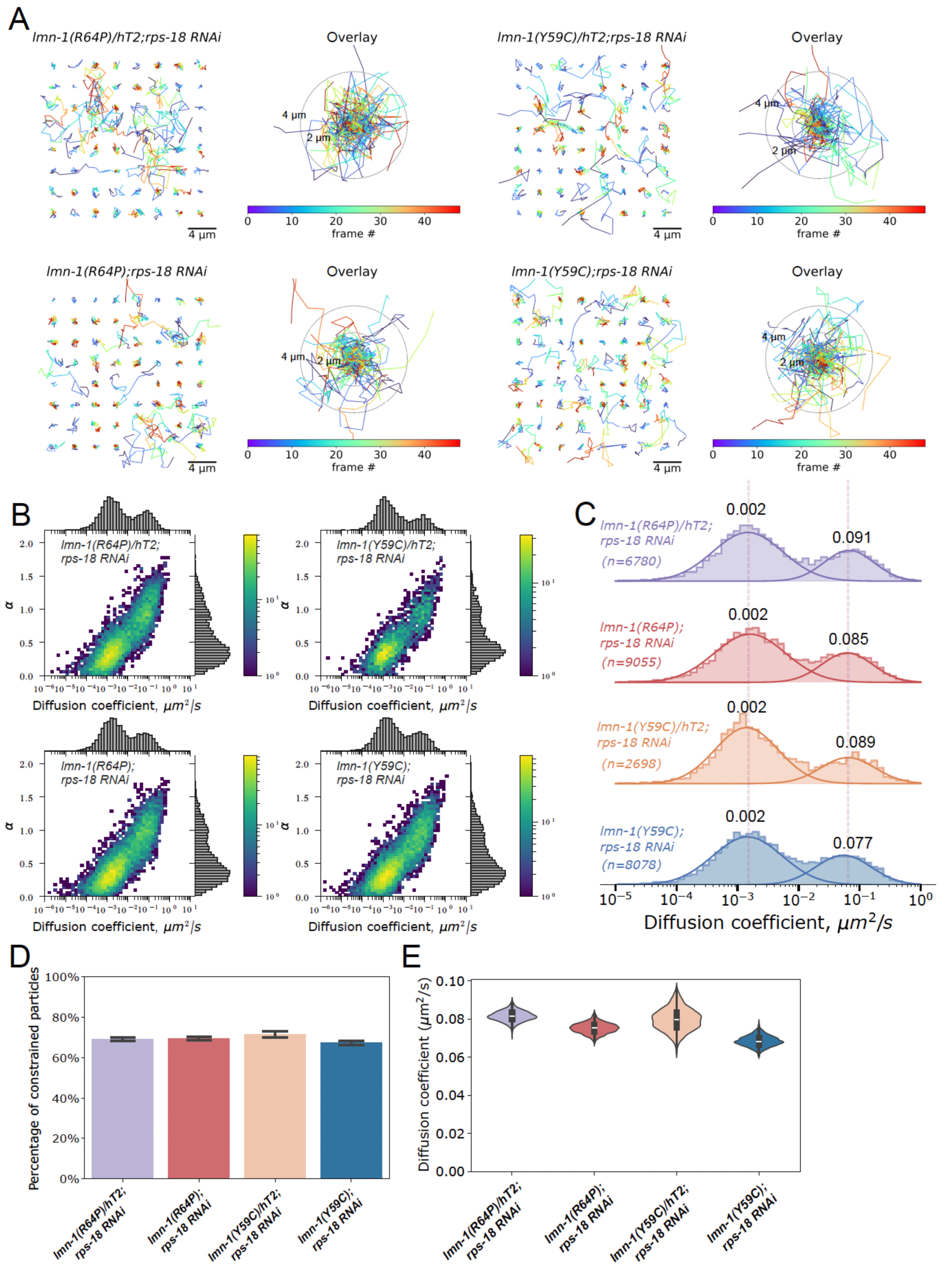
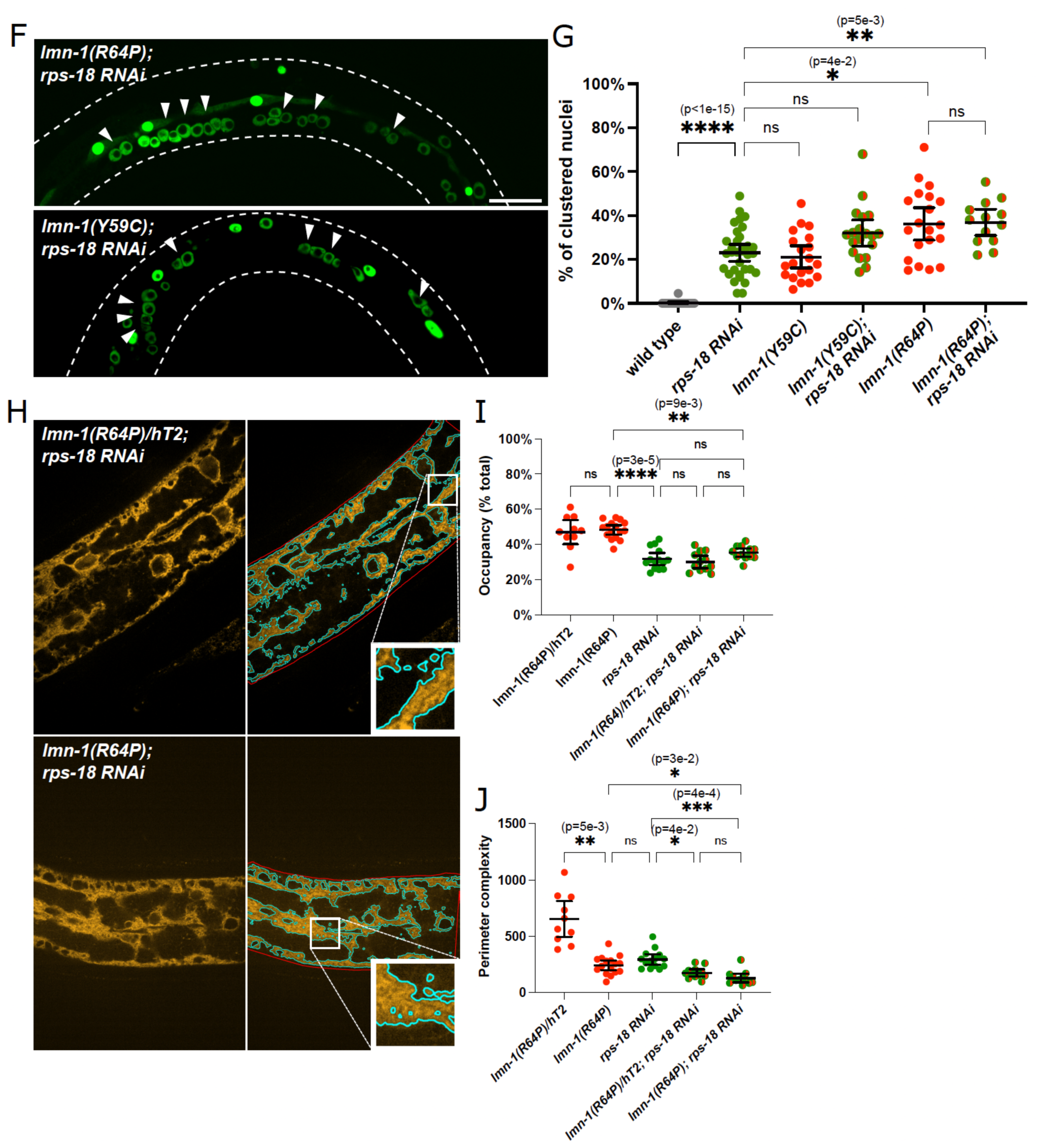
LMN-1 and ribosomes act in the same pathway to promote cytoplasmic organization. **A)** Representative particle trajectories color-coded by frame number for GEMs tracked in the intestine of the indicated strains. Scale bar: 4 μm. Concentric circles indicate 2 μm and 4 μm radii from trajectory origin. **B)** 2D probability density histograms correlating GEM D_eff_ with α in the intestinal cytoplasm for the indicated strains at day 1 of adulthood. Marginal distributions shown as frequency histograms on top and right axes. Color scale indicates particle count per bin. **C)** GMM analysis of GEM *D_eff_* in the intestine of the indicated strains at day 1 of adulthood. Dotted lines indicate the center of each distribution (purple: *lmn-1(R64P)/hT2;rps-18(RNAi)*, and dark red: *lmn-1(R64P); rps-18(RNAi)*). **D-E)** Bootstrap analysis of GMM parameters comparing GEM diffusion in the intestine of the indicated strains at day 1 of adulthood. The percentage of constrained GEMs (D) and mean ± SD *D_eff_* values of unconstrained GEMs (E) are shown. **F)** Representative spinning disc confocal images of GFP-labeled hypodermal nuclei in day 1 adults of the indicated strains. Arrowheads indicate defective nuclear anchoring; dotted lines outline worm boundaries. Scale bar: 10 μm. **G)** Nuclear positioning defects quantified as percentage of touching nuclei per worm in day 1 adults of the indicated strains. Left to right: n1=20, n2=33, n3=20, n4=19, n5=20, n6=13. Welch’s ANOVA test: W(5, 43) = 115, p < 0.001. **H)** Representative spinning disc confocal images of hypodermal ER organization in the indicated strains at day 1 of adulthood. Left panels: ER morphology visualized by mKate2::TRAM-1 fluorescence. Right panels: Thresholded images showing ER outline (cyan) and hypodermal boundary (red). Scale bar: 10 μm. Zoom-in insets are magnified views of the boxed regions. **I-J**) ER network analysis showing ER occupancy percentage (I) (n=10, 16, 14, 12, 12 from left to right), Kruskal-Wallis test: H(4)=40, p <0.001, and perimeter complexity (J) (n=10, 16, 14, 12, 12 from left to right), Kruskal-Wallis test: H(4)=44, p<0.001, across genotypes. Statistical significance is indicated by asterisks (****: p<0.0001, ***: p≤0.001,**: p≤0.01, *: p≤0.05, ns: p>0.05) with exact p-values shown above each comparison.

Nuclear positioning assays revealed similar epistatic patterns. While *rps-18(RNAi)* slightly enhanced nuclear clustering defects in *lmn-1(Y59C)* mutants, it produced no additional defect in *lmn-1(R64P)* backgrounds (**Figs. 3H-I**). This consistent pattern across two independent cellular phenotypes strengthened the evidence for pathway convergence and indicated that genetic interactions were not specific to cytoplasmic crowding effects alone. As predicted by the pathway model, the ER morphology parameters in *rps-18(RNAi); lmn-1(R64P)* double mutant animals remained comparable to *rps-18(RNAi)* heterozygous controls (**Figs. 3J-L**), completing the epistatic pattern across all three cellular organization phenotypes we examined. Thus, across all three phenotypes, LMN-1 and ribosomes act in a common pathway controlling cellular organization.

### LMN-1 regulates the nucleolar-ribosome axis

Having demonstrated that lamins and ribosomes function in the shared pathway to regulate cellular organization, we sought to determine the mechanistic relationship between these factors. Ribosomes are synthesized and assembled within the nucleolus before export to the cytoplasm^12^. Previous studies in mammalian tissue culture cells showed that lamins play important roles in nucleolar function^24,25^, but the effects of EDMD-associated lamin mutations on nucleolar organization and ribosome production in intact animal tissues remained unexplored. We hypothesized that lamin dysfunction disrupts cellular organization by interfering with ribosome biogenesis in the nucleolus.

To test this hypothesis, we examined nucleolar organization in *lmn-1* mutant animals using FIB-1::GFP, a fluorescently tagged fusion of the *C. elegans* fibrillarin ortholog, an essential nucleolar protein^26^. Fibrillarin is a well-established marker of nucleolar activity whose levels correlate directly with ribosome biogenesis capacity across multiple species^12,27–29^. Live imaging of nucleoli in the hypodermal syncytium allowed us to quantify both nucleolar morphology and FIB-1 protein abundance (**Figs. 4A**). While *lmn-1(R64P)* mutants showed nucleolar sizes comparable to balanced heterozygous *lmn-1(R64P)/hT2* controls (**Figs. 4B**), they exhibited significant reduction in FIB-1::GFP fluorescence intensity (**Figs. 4C**). The decrease in FIB-1::GFP intensity within nucleoli of normal size suggests that the *lmn-1(R64P)* mutation may impair the recruitment, retention, or stability of factors involved in ribosome biogenesis.

**Figure 4.**
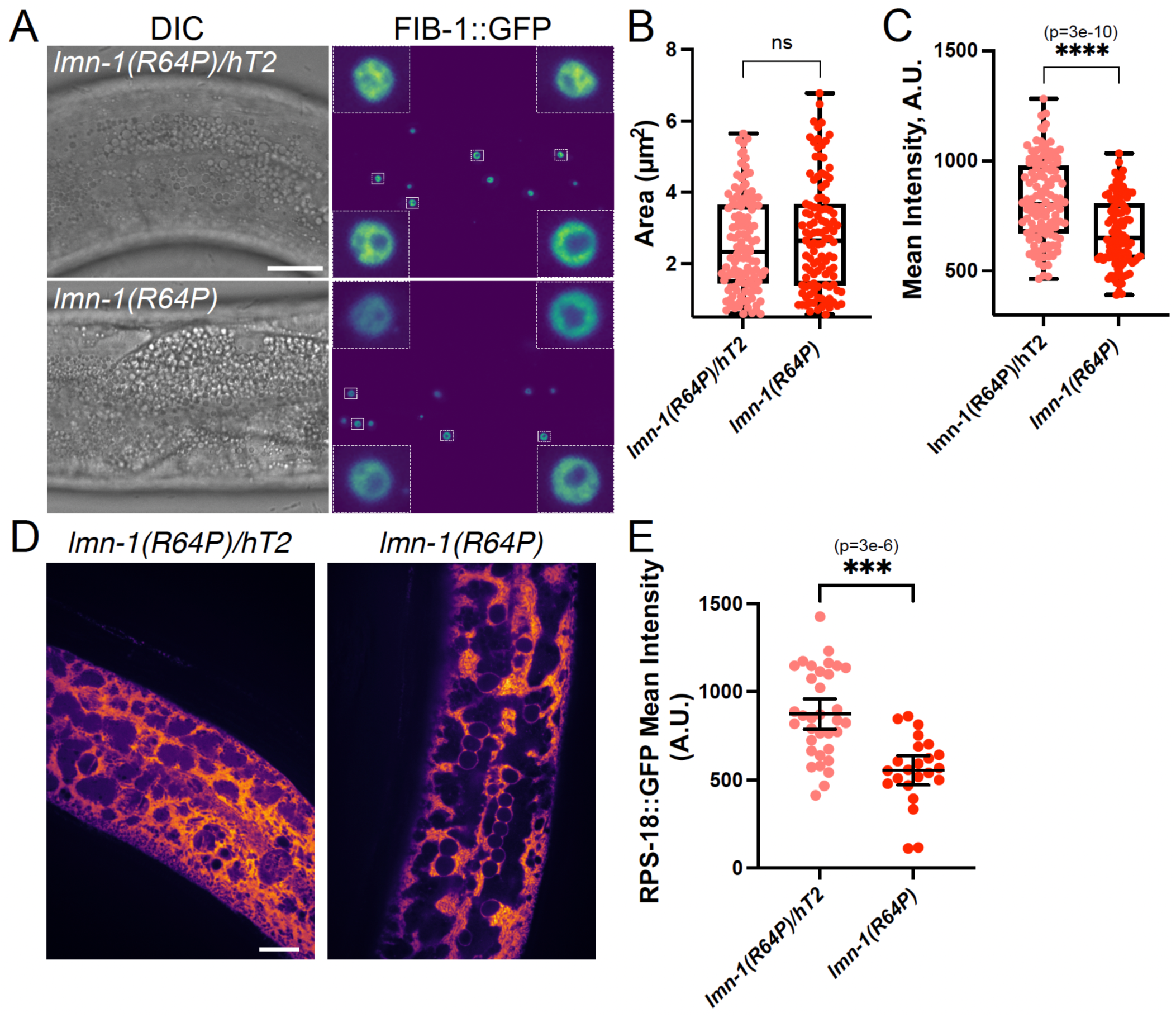
*lmn-1(R64P)* mutants show reduced nucleolar FIB-1 density and ribosome abundance. **A)** Representative spinning-disk confocal images (right) of FIB-1::GFP in the hypodermis of the indicated strains at day 1 of adulthood, alongside corresponding differential interference contrast (DIC) images (left). Insets depict magnified views of boxed nucleoli. Scale bar: 10 µm. Fluorescence intensity is shown using the viridis colormap. **B)** Quantification of individual nucleoli as visualized with FIB-1::GFP fluorescence. Each dot represents an individual nucleolus from a total of 9 worms per strain. Statistical significance was assessed by two-tailed Welch’s t-test. Welch’s test: t(217.5)=1.274, p >0.05. **C)** Quantification of mean FIB-1::GFP fluorescence intensity per nucleolus from a total of 9 worms per strain (left to right, n=126, 111). Welch’s test: t(234.2) = 6.574, p < 0.001. **D)** Representative spinning disc confocal images of RPS-18::GFP expressed in the hypodermis of the indicated strains at day 1 of adulthood. Scale bar: 10 μm. **E)** Quantification of mean RPS-18::GFP fluorescence intensity per hypodermal image across the indicated genotypes (left to right: n=34, 23). Welch’s test: t(53.73)=5.442, p<0.001. Statistical significance is indicated by asterisks (****: p<0.0001, ***: p≤0.001,**: p≤0.01, *: p≤0.05, ns: p>0.05) with exact p-values shown above each comparison.

To determine whether reduced nucleolar FIB-1::GFP intensity leads to decreased ribosome production, we measured ribosome levels using RPS-18::GFP^10,30^. We observed significant decreases in hypodermal RPS-18::GFP fluorescence intensity in *lmn-1(R64P)* mutants compared to heterozygous *lmn-1(R64P)/hT2* controls (**Figs. 4D-E**). The coordinated reduction in FIB-1 (nucleolar assembly factor) and RPS-18 (ribosomal component) indicates that lamin dysfunction disrupts ribosomal biogenesis. These findings demonstrate a mechanism whereby lamin mutations impair nucleolar function, leading to reduced ribosome production and subsequent disruption of cytoplasmic organization. This nucleolar-ribosome axis provides a molecular explanation for how nuclear lamina defects propagate throughout the cell to cause the broad cellular disorganization characteristic of EDMD.

### Cytoplasmic crowding defects are specific to EDMD-associated nuclear envelope proteins

To determine whether cytoplasmic organization defects were specific to EDMD-associated proteins or represent a general consequence of nuclear envelope disruption, we examined how various nuclear envelope protein mutations affect cytoplasmic biophysical properties using GEM particle tracking. In humans, loss of emerin alone causes X-linked EDMD^1^ However, in *C. elegans*, the emerin ortholog EMR-1 functions redundantly with another LEM-domain protein, LEM-2, during early embryogenesis^31,32^. Previous studies showed that co-depletion of *emr-1* and *lem-2* recapitulate nuclear envelope defects and EDMD-like phenotypes observed in LMN-1-depleted animals^17,31,33^. We found that single *lem-2(tm1582)* or *emr-1(gk119)* mutations produced minimal disruption to GEM diffusion patterns (**Figs. 5A-F, S3**). However, *emr-1(gk119); lem-2(RNAi)* double mutants led to disruption of cytoplasmic crowding (**Figs. 5G-I**). *emr-1(gk119); lem-2(RNAi)* animals had marked increased GEM diffusion coefficients, mirroring lamin mutants, demonstrating that combined loss of functionally redundant LEM-domain proteins resembles lamin dysfunction.

**Figure 5.**
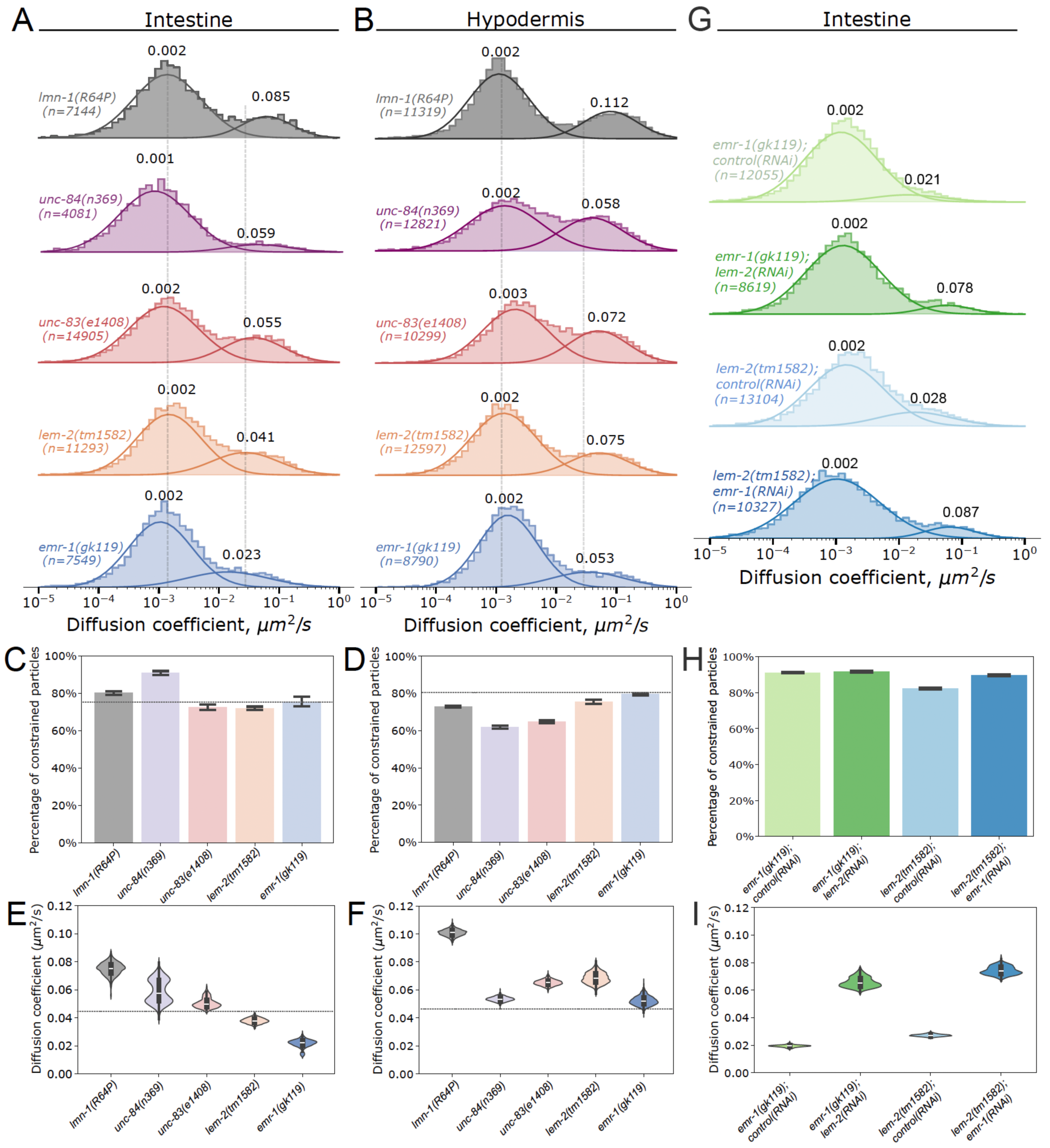
Combined loss of LEM-domain proteins mimics lamin dysfunction and disrupts cytoplasmic crowding. **A-B, G)** Gaussian mixture model (GMM) analysis of GEM *D_eff_* in the intestine (A) or hypodermis (B, G) of the indicated strains at day 1 of adulthood. Gray dotted lines mark the population center of lmn-1(R64P)/hT2 animals as a reference to guide the eye. C-F, H-I) Bootstrap analysis of GMM parameters comparing GEM diffusion in the intestine (C, E, H, I) or hypodermis (D, F). Shown are the percentage of constrained GEMs (C–D, H) and D_eff_ values of unconstrained GEMs (E–F, I). Gray dotted lines in each plot indicate the mean value of lmn-1(R64P)/hT2 for reference. Data are presented as mean ± SD.

We also tested canonical LINC complex components, including the inner nuclear membrane Sad1/UNC-84 (SUN) domain protein UNC-84 and the outer nuclear membrane partner, the KASH protein UNC-83^34–36^. These proteins are core constituents of the LINC complex that mediates nuclear-cytoskeletal coupling and interact with lamin^5^. In mammals, LINC complex components such as SUN1, SUN2, nesprin-1, and nesprin-2 have also been implicated in EDMD^37^. However, animals carrying null mutations *unc-84(n369)* or *unc-83(e1408)* displayed only minimal alterations in cytoplasmic biophysical properties in both intestine and hypodermis (**Figs. 5A-F, S3**).

These results indicate that disruption of individual nuclear envelope components does not substantially affect cytoplasmic organization. Across all single mutants, the percentage of constrained particles remained largely unchanged (**Figs. 5C-D, S3**), and diffusion coefficients of unconstrained particles showed only modest variations (**Figs. 5E-F**). Thus, cytoplasmic crowding defects are not a general consequence of nuclear envelope dysfunction but instead represent a specific hallmark of EDMD-associated protein disruption. The pronounced effect of combined emerin/LEM-domain perturbation, contrasted with the minimal impact of other nuclear envelope components loss, highlights the distinctive requirement for EDMD-linked proteins in maintaining cytoplasmic biophysical properties.

## DISCUSSION

This study uncovers a nuclear lamina-nucleolar-ribosomal axis through which defects in nuclear lamins propagate to the cytoplasm, disrupting cellular architecture. While previous studies identified connections between lamins and nucleolar function in cultured mammalian cells^24,25^, our work demonstrates how this relationship controls cytoplasmic biophysical properties and cellular organization in intact tissues. Using *C. elegans*, we show that EDMD-associated lamin variants reduce mesoscale cytoplasmic crowding, misposition nuclei, and collapse ER network complexity through effects on nucleolar organization and ribosome production (**Fig. 6**).

**Figure 6.**
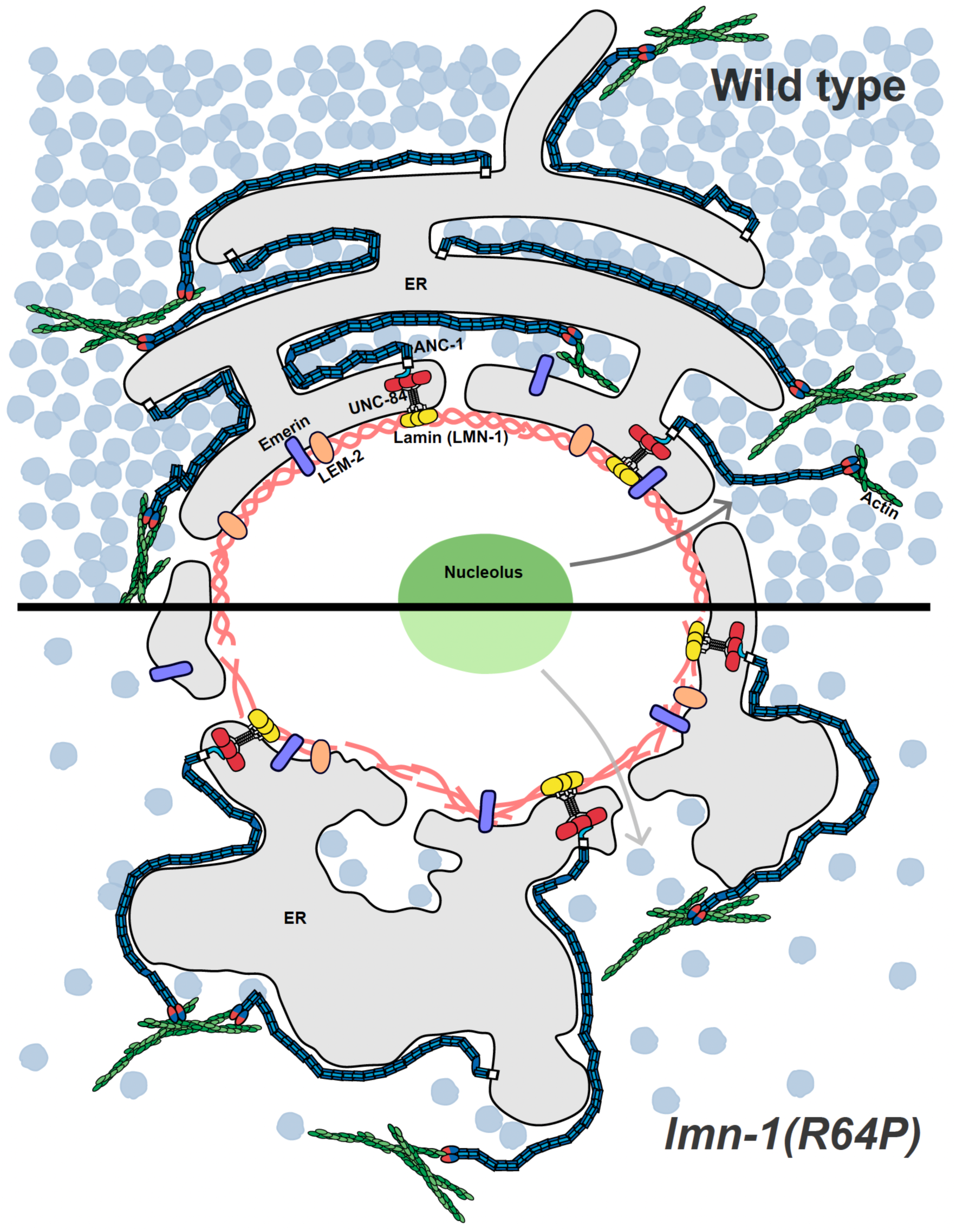
Nuclear lamina maintains cellular organization through a nucleolar-ribosome axis. Top (wild type): An intact nuclear lamina preserves nucleolar integrity, supporting ribosome biogenesis. High ribosome abundance sustains cytoplasmic macromolecular crowding, stabilizing ER network morphology and cellular organization. Bottom (lmn-1(R64P)): Lamina dysfunction disrupts nucleolar organization, reduces ribosome production, and lowers cytoplasmic crowding. This loss of crowding leads to ER network collapse, elevated cytoplasmic diffusion, and global cellular disorganization.

### Integration with existing laminopathy models

Two prevailing models have historically shaped explanations for how lamin mutations drive pathology: 1) lamina defects weaken nuclear architecture and mechanotransduction, rendering muscle nuclei vulnerable to mechanical stress and aberrant signaling; and 2) altered chromosome organization disrupts gene regulation, impairing tissue-specific gene activation^8^. Our nuclear lamina-nucleolar-ribosome framework complements these models by revealing how nuclear defects translate into widespread cytoplasmic disorganization characteristic of EDMD.

While mechanotransduction models emphasize nuclear membrane integrity and force transmission, and the gene regulation models highlight transcriptional control, we propose that the lamina-nucleolar-ribosome axis may function downstream of both mechanisms. Nucleolar dysfunction could result from either compromised nuclear mechanics or altered ribosomal RNA transcription^25,38–42^. In support of this idea, we show that EDMD-linked *lmn-1* mutations reduce ribosome biogenesis and in turn impair cytoplasmic biophysical properties, thereby linking nuclear lamina defects directly to cytoplasmic disorganization. Importantly, our model incorporates cytoplasmic phenotypes that earlier frameworks have not adequately addresses, including altered macromolecular crowding, disrupted ER architecture and organelle mispositioning. These observations resonate with recent work demonstrating that mesoscale condensates represent a fundamental organizing principle of the cytoplasm reinforcing the view that cytoplasmic organization emerges from active, regulated processes vulnerable to perturbation by nuclear envelope dysfunction^43^.

### Evolutionary conservation and clinical relevance

The nucleolar-ribosome axis is highly conserved across species^44,45^, suggesting our findings in *C. elegans* are likely relevant to human disease. Mammalian lamins directly interact with nucleolar components and lamin mutations in patient-derived cells show nucleolar abnormalities^24,25^, resembling our observations. The conservation of ribosome-dependent cytoplasmic crowding from *E. coli* to mammals^9,46,47^ further supports the translational relevance of our findings.

This conservation has important clinical implications. If nucleolar dysfunction and ribosome depletion were central to EDMD pathogenesis, therapeutic strategies could target multiple points in this pathway: enhancing ribosome biogenesis, modulating mTOR signaling to restore ribosome production, or using crowding agents to compensate for reduced ribosome levels. Such approaches could potentially benefit patients with mutations in either *LMNA* or *EMD*, providing broader therapeutic options than mutation-specific strategies.

### Tissue specificity and disease progression

The variable tissue sensitivity we observed between intestine and hypodermis provides insights into EDMD’s characteristic tissue-specific pathology. Skeletal and cardiac muscle may be particularly vulnerable because they have high mechanical demands that require robust cytoplasmic organization, extensive ER networks for calcium handling, and precise nuclear positioning^3,22^. Tissues with lower mechanical stress of different metabolic demands might tolerate reduced cytoplasmic crowding more effectively, explaining the selective tissue involvement in EDMD.

### Mechanistic insights from genetic interactions

The phenotypic parallels between EDMD-associated lamin mutants and ribosome depletion are striking. Both conditions cause nuclear positioning and ER morphology defects. Severity differences between R64P and Y59C mutants match the known severity of polymerization defects caused by the R64P mutation compared to Y59C^13,17^, and epistasis analyses demonstrate that R64P operates in the same genetic pathway as ribosomes for regulating cytoplasmic organization. In contrast, the giant KASH protein ANC-1 functions in a second pathway by scaffolding the ER and constraining the cytoplasm^10^. Together, quantification of nucleolar FIB-1 and ribosome levels positions lamin defects upstream of ribosome biogenesis, clarifying how nuclear lamina mutations trigger cytoplasmic disorganization in EDMD.

### Resolving the mTOR paradox

A paradox emerges when considering the hyperactivation of mTORC1 signaling in laminopathies^48^. Given mTORC1’s role as a master regulator promoting ribosome biogenesis^49^, how do lamin mutations simultaneously cause mTOR hyperactivation and ribosome depletion? This apparent contradiction suggests that additional regulatory mechanisms must override mTOR’s pro-ribosomal signals in laminopathies. Lamins contribute to nucleolar structural integrity and stress response regulation^25,50,51^, and pathogenic variants frequently trigger an ER stress response that suppresses rRNA transcription^52^. Thus, despite elevated mTOR signaling, the combination of nucleolar dysfunction and ER stress may reduce overall ribosome production through stress-activated pathways that override growth-promoting signals. This model reconciles the apparent contradiction and suggests that therapeutic interventions targeting stress responses together with mTOR might be more effective.

### Technical advances and broader applications

The development of GEM-based nanorheology for intact tissues represents a significant methodological advance with applications beyond laminopathy research. Similar nuclear GEM approaches were recently applied in mammalian cells to probe nuclear mechanical properties^53^. By applying tissue-specific GEM expression, we learned that *C. elegans* maintains distinct cytoplasmic biophysical properties: both crowded and constrained. Our study showed that EDMD-associated lamin mutations selectively reduce the crowding of unconstrained cytoplasmic particles, suggesting a role in regulating the concentration or organization of crowding agents rather than structural barriers. Given that ribosome concentration is a primary determinant of cytoplasmic crowding in multiple species^10,46,47^, these results strongly suggest that lamin mutations impair ribosome levels, distribution, or assembly. This approach could be applied to study cytoplasmic biophysics in other diseases involving protein aggregation, metabolic disorders affecting ribosome function, or aging-related changes in cellular organization. The ability to quantify cytoplasmic crowding in living animals provides a new biomarker for disease severity and therapeutic response that was previously unavailable.

### Therapeutic implications and future directions

Our finding that *emr-1* and *lem-2* function redundantly to regulate cytoplasmic crowding is relevant because these encode LEM domain proteins, including the *C. elegans* emerin ortholog^32^,and mutations in human emerin cause X-linked EDMD^1^. The *emr-1; lem-2* double mutants phenocopy lamin mutations in *C. elegans* embryonic mitosis and viability, further supporting that they function in a common pathway^32^. Thus, both major EDMD-associated proteins, lamin and emerin, contribute to regulating cytoplasmic biophysical properties.

The partial functional redundancy between *emr-1* and *lem-2* may explain why EDMD pathology is variable and tissue-specific, as functional compensation between nuclear envelope proteins could mask defects in most tissues. The identification of the lamina-nucleolar-ribosome axis as a common pathway affected in both lamin and emerin associated forms of EDMD suggests potential therapeutic opportunities. GEM-based nanorheology could serve as a useful tool for monitoring cellular organization in preclinical studies.

## Conclusion

This work establishes a mechanistic link between nuclear lamina defects, nucleolar function, ribosome biogenesis, and cytoplasmic biophysical properties, revealing how nuclear lamina dysfunction propagates to disrupt cellular organization throughout the cytoplasm. This represents a previously uncharacterized mechanism of laminopathy pathogenesis that connects nuclear architecture to fundamental cellular biophysics. The development and application of *in vivo* quantitative GEM-based nanorheology represents a significant technical advance, enabling quantitative analysis of cytoplasmic biophysical properties in intact animal tissues. Our findings demonstrate that both major EDMD-associated proteins, lamin and emerin, regulate cytoplasmic organization by maintaining cytoplasmic biophysical properties, offering new insights into the molecular basis of EDMD and potential therapeutic targets for treating laminopathies.

## Methods

### *C. elegans* strains and maintenance

All *C. elegans* strains used in this study are listed in Supplemental Table S1. Worms were maintained on standard nematode growth medium (NGM) agar plates seeded with *Escherichia coli* strain OP50 at 23° C^54^. For all experiments, synchronized animals were cultured to young adult stage (24 hours post-L4) under non-starved conditions. Experimental animals were maintained on fresh plates for a minimum of two generations prior to experimentation to ensure consistent physiological conditions.

### RNA interference

Gene knockdown was performed using standard feeding RNAi methodology^55,56^. Bacterial strains harboring specific RNAi clones were obtained from the Ahringer RNAi library (Source Bioscience, Nottingham, UK) and verified by Sanger sequencing prior to use. Individual bacterial colonies were picked from fresh plates and grown overnight at 37° C in 1 mL LB medium supplemented with 100 μg/mL carbenicillin in 15 mL snap-cap tubes on a rotary shaker. Cultures were diluted 1:40 in 2 mL fresh LB-carbenicillin medium and grown for 3 hours at 37° C. For dsRNA induction, 2 mL of pre-warmed LB medium containing 100 μg/mL carbenicillin and 1 mM isopropyl β-D-1-thiogalactopyranoside (IPTG) was added to achieve a final IPTG concentration of 0.5 mM, followed by incubation for 3-4 hours at 37° C. Induced bacterial cultures were spotted onto NGM agar plates containing 25 μg/mL carbenicillin and 1 mM IPTG. Synchronized L2-L3 stage larvae were transferred to RNAi feeding plates and maintained at 23° C for 48 hours until reaching young adult stage. To achieve optimal GEM expression levels for single-particle tracking in hypodermal tissue, L3 stage worms expressing GEM-EGFP were fed bacteria expressing dsRNA targeting *gfp* for 48 hours until egg-laying commenced^10^. L1 progeny were then transferred to standard NGM plates with OP50 bacteria and cultured until imaging age. This protocol reduced GEM concentration to appropriate levels for tracking analysis.

### GEM detection and motion analysis

GEMs were imaged as previously reported^10^. In brief, live imaging was performed using a Nikon Ti2 inverted microscope (Nikon Instruments, Melville, NY) equipped with a Yokogawa CSU-X1 spinning disk confocal scanner unit (Yokogawa Electric Corporation, Tokyo, Japan) and a Hamamatsu ORCA-Flash4.0 LT3 Digital sCMOS camera (Hamamatsu Photonics, Shizuoka, Japan). Images were acquired using a Nikon Plan Apo λ 100× oil immersion objective (N.A. 1.45) at 1060 × 1568 pixels resolution (0.065 μm/pixel). GEMs were visualized using 488 nm laser excitation at 100% power with 20 ms exposure time per frame in continuous acquisition mode (no inter-frame interval). Time-series data were collected at 50 Hz acquisition rate for motion analysis. GEM particle detection and tracking were performed using published methods^10^. Briefly, trajectories shorter than 10 frames (0.2 seconds) were excluded from mean squared displacement (MSD) analysis to ensure statistical reliability. For each valid trajectory, the effective diffusion coefficient (*D_eff_*) was calculated by linear fitting of the first 5 data points (100 ms total duration) of the MSD curve. Two-component Gaussian mixture models were fitted to *D_eff_* distributions to resolve sub-populations of constrained and unconstrained particles. Parameter distributions and confidence intervals were estimated using bootstrap resampling with 1,000 iterations^10^.

### ER imaging and morphological analysis

ER networks were visualized using the mKate2::TRAM-1 fluorescent marker (strain UD756, Table S1). Images were acquired using the Nikon Ti2 microscope system described above with 561 nm laser excitation at 50% power, 100 ms exposure time, and identical objective and camera configurations. ER occupancy and complexity analyses were carried out as previously described^10,23^. Briefly, images underwent background subtraction and noise reduction before binary segmentation using Otsu’s thresholding method. Two morphometric parameters were quantified: (1) ER occupancy, defined as the ratio of ER-positive area to total cellular area, and (2) ER network complexity, calculated as the perimeter-to-area ratio (P²/4πA), where P represents the total perimeter and A represents the total area of ER-positive regions.

### Nuclear positioning measurements

Hypodermal nuclei were visualized using nuclear-localized GFP expressed from the integrated transgene *ycIs10[p_col-10_::nls::*GFP*::lacZ]*^20^. Young adult worms were cultured on either standard NGM plates with OP50 *E. coli* (controls) or RNAi feeding plates with HT115 *E. coli* expressing target dsRNA. Microscope slides were prepared as introduced previously^20^. Nuclear positioning was assessed using an Andor BC43 microscope with 488 nm laser excitation and a 60×/1.42 oil immersion objective. To ensure systematic counting, the anus and pharynx of each worm were positioned in the same focal plane. Nuclear clustering was quantified by counting GFP-positive hypodermal nuclei in direct contact with neighboring nuclei^20,21^. Contacts along the perpendicular axis were excluded from analysis as the nuclear marker does not distinguish seam cell nuclei from hyp7 nuclear clusters. The percentage of clustered nuclei was calculated as the number of nuclei in contact divided by the total number of nuclei counted per animal.

### Nucleolar measurements

Nucleoli in both intestinal and hypodermal tissues were visualized using FIB-1::GFP^57^, which is expressed pan-tissue wise throughout the organism. Worms were synchronized to L4 stage and imaged 24 hours later as young adults. Animals were immobilized using 10 μM tetramisole in M9 buffer on 2% agarose pads using standard procedures^20^. Confocal microscopy on BC43 was performed using an Andor BC43 microscope with 488 nm laser excitation (3.0% laser power, 100 ms exposure time) and a 60×/1.42 oil immersion objective. Brightfield imaging was performed concurrently for tissue localization. Nucleolar segmentation was performed using Otsu’s automatic thresholding method^58^ in FIJI/ImageJ software, with size filters applied (0.5 μm² to infinity) to exclude artifacts. Regions of interest encompassing all identified nucleolar objects were generated and applied to original raw images for fluorescence intensity measurements. Mean fluorescence intensity was measured for each segmented nucleolus.

### Ribosome visualization and analysis

Ribosomes were visualized using an endogenously tagged RPS-18 protein created through CRISPR/Cas9 genome editing to generate the allele *yc113[*GFP11*::rps-18] IV*^10^. The complementary GFP_1-10_ construct was expressed in hypodermal tissue using the transgene JuEx5375[*p_col-19_*::GFP_(1-10)_+ *p_ttx-3_*::RFP]^30^, enabling reconstitution of functional GFP fluorescence specifically in the hypodermis. Young adult worms were immobilized using 10 μM tetramisole in M9 buffer on 2% agarose pads as described above. Images were acquired using the above-described Nikon Ti2 microscope system with a 100× oil immersion objective (N.A. 1.45) at 1060 × 1568-pixel resolution (0.065 μm/pixel). Ribosome fluorescence was captured using 488 nm laser excitation at 30% power with 50 ms exposure time. Hypodermal cell boundaries were manually delineated to define regions of interest for fluorescence intensity measurements. Average ribosome fluorescence intensity was quantified within the hypodermal syncytium.

### Statistical analyses

All statistical analyses were performed using Prism GraphPad software (GraphPad Software, Inc.). Sample sizes (n), degree of freedom and exact p-values are reported in figure legends for each experiment. Nuclear positioning defects were analyzed using Welch’s ANOVA to account for unequal variances, followed by Dunnett’s T3 post hoc test for multiple comparisons. ER morphological defects were analyzed using Kruskal-Wallis followed by Dunnett’s T3 post hoc test for multiple comparisons. For nucleolus and ribosome quantification, two-tailed Welch’s test was performed. All Statistical significance is indicated by asterisks (****: p<0.0001, ***: p≤0.001,**: p≤0.01, *: p≤0.05, ns: p>0.05) with exact p-values shown above each comparison in each legend.

---

**Table S1.** Strain list.

| Strain Name | Genotype | References |
| --- | --- | --- |
| UD803 | *ycSi4*[pSL848; *pges-1::40nmGEM::eGFP::unc-54 3'UTR*] IV | <sup>10</sup> |
| UD838 | *ycSi5*[pSL850; *psemo-1::40nmGEM::eGFP::unc-54 3'UTR*] IV | <sup>10</sup> |
| UD1055 | *lmn-1*(*yc96*[Y59C]) *I/hT2* [*bli-4*(e937) *let*(q782) *qls48*[*P<sub>myo-2</sub>::gfp*; *P<sub>pes-10</sub>::gfp*; *P<sub>ges-1</sub>::gfp*]] (*I*; *III*); *ycSi4* IV | This study |
| UD1054 | *lmn-1*(*yc96*[Y59C]) *I/hT2* (*I*; *III*); *ycSi5* IV | This study |
| UD1043 | *lmn-1*(*yc105*[R64P]) *I/hT2* [*bli-4*(e937) *let*(q782) *qls48*[*P<sub>myo-2</sub>::gfp*; *P<sub>pes-10</sub>::gfp*; *P<sub>ges-1</sub>::gfp*]] (*I*; *III*); *ycSi4* IV | This study |
| UD916 | *lmn-1*(*yc105*[R64P]) *I/hT2* (*I*; *III*); *ycSi5* IV | This study |
| UD1031 | *lmn-1*(*tgx535*[R204W]) *I*; *ycSi4* IV | This study |
| UD1038 | *lmn-1*(*tgx535*[R204W]) *I*; *ycSi5* IV | This study |
| UD1042 | *lmn-1*(*yc105*) *I/hT2* (*I*,*III*); *ycSi2* [pSL845 *Ppy37A1B.5::mKate2::tram-1*] IV | This study |
| COP262 | *unc-119*(ed3) *III*; *knuSi221* [*P<sub>fib-1</sub>::fib-1::eGFP::fib-1 3'UTR+unc-119(+)*] *II* | <sup>57</sup> |
| UD1079 | *knuSi221* *II*; *lmn-1*(*yc105*[R64P]) *I/hT2* (*I*; *III*); *ycSi2* IV | This study |
| UD863 | *yc113* [*GFP<sub>11</sub>::rps-18*] IV; *ycSi2* IV; *juEx5375* [*P<sub>col-19</sub>::GFP<sub>1-10</sub>+ P<sub>ttx-3</sub>::RFP*] | <sup>10</sup> |
| UD1099 | *yc113* IV; *juEx5375*; *lmn-1*(*yc105*) *I/hT2* (*I*,*III*); *ycSi2* IV | This study |
| UD883 | *lmn-1* (*yc96*) *I/hT2* (*I*,*III*); *ycls10*[*P<sub>col-10</sub>::nls::gfp::lacZ*] V, *him-8* (e1489) IV | <sup>15</sup> |
| UD895 | *lmn-1* (*yc105*) *I/hT2* (*I*,*III*); *ycls10* V, *him-8* (e1489) IV | <sup>15</sup> |
| UD908 | *lmn-1*(*tgx535*[R204W]) *I*; *ycls10* V, *him-8*(e1489) IV | <sup>15</sup> |
| UD1041 | *emr-1*(*gk119*) *I*; *ycSi4* IV | This study |
| UD1044 | *emr-1*(*gk119*) *I*; *ycSi5* IV | This study |
| UD1046 | *lem-2*(*tm1582*) *II*; *ycSi4* IV | This study |
| UD1045 | *lem-2*(*tm1582*) *II*; *ycSi5* IV | This study |
| UD977 | *unc-84*(n369) X; *ycSi4* IV | This study |
| UD875 | *unc-84*(n369) X; *ycSi5* IV | This study |
| UD1018 | *unc-83*(e1408) V; *ycSi4* IV | This study |
| UD1019 | *unc-83*(e1408) V; *ycSi5* IV | This study |

**Figure S1.**
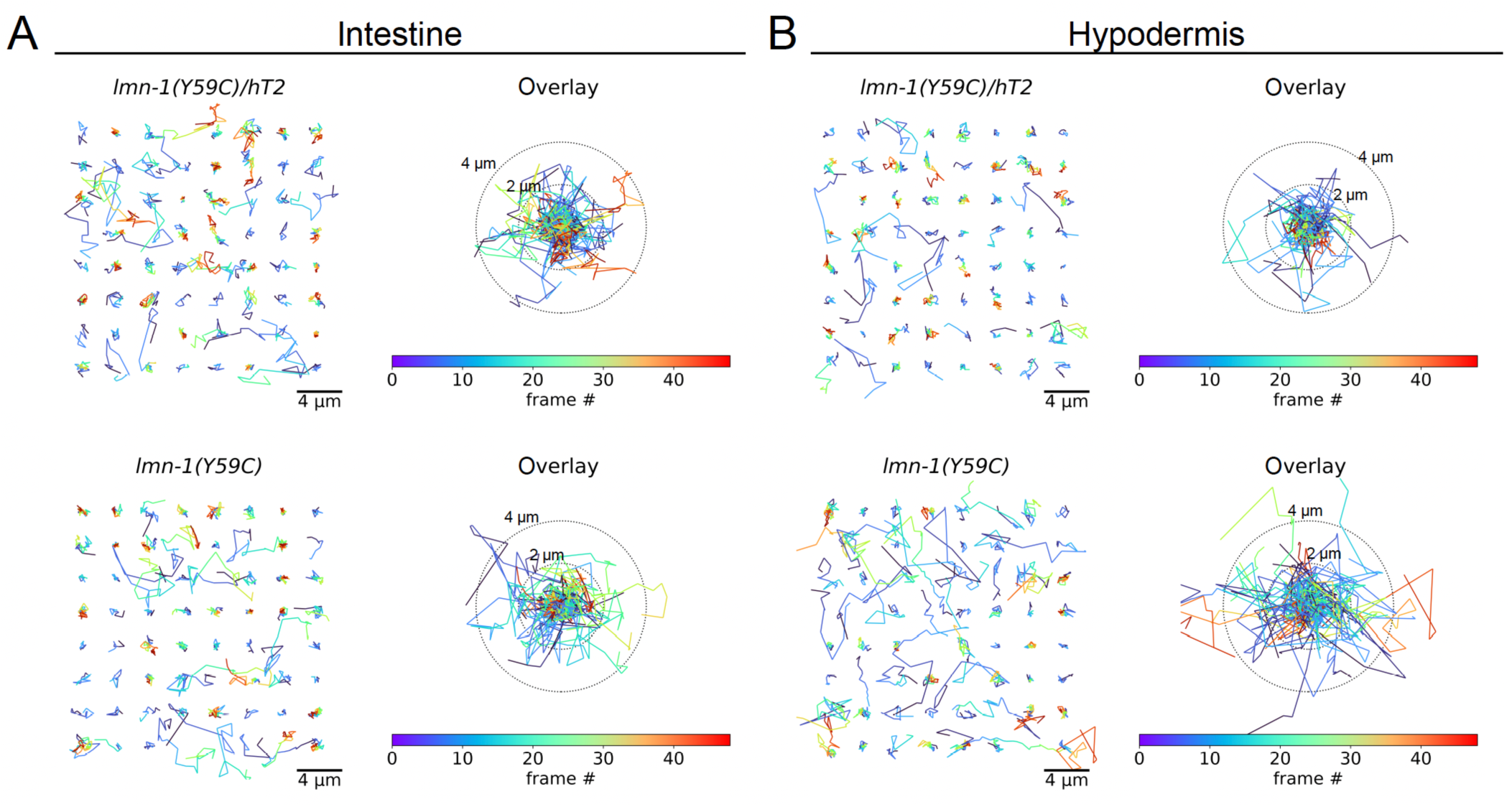
Homozygous *lmn-1(Y59C)* mutants exhibit increased GEM mobility in the hypodermis but show minimal changes in the intestine. Representative GEM trajectories color-coded by frame number are shown for (A) intestine and (B) hypodermis of the indicated strains. Scale bar: 4 µm. Concentric circles denote 2 µm and 4 µm radii from the trajectory origin.

**Figure S2.**
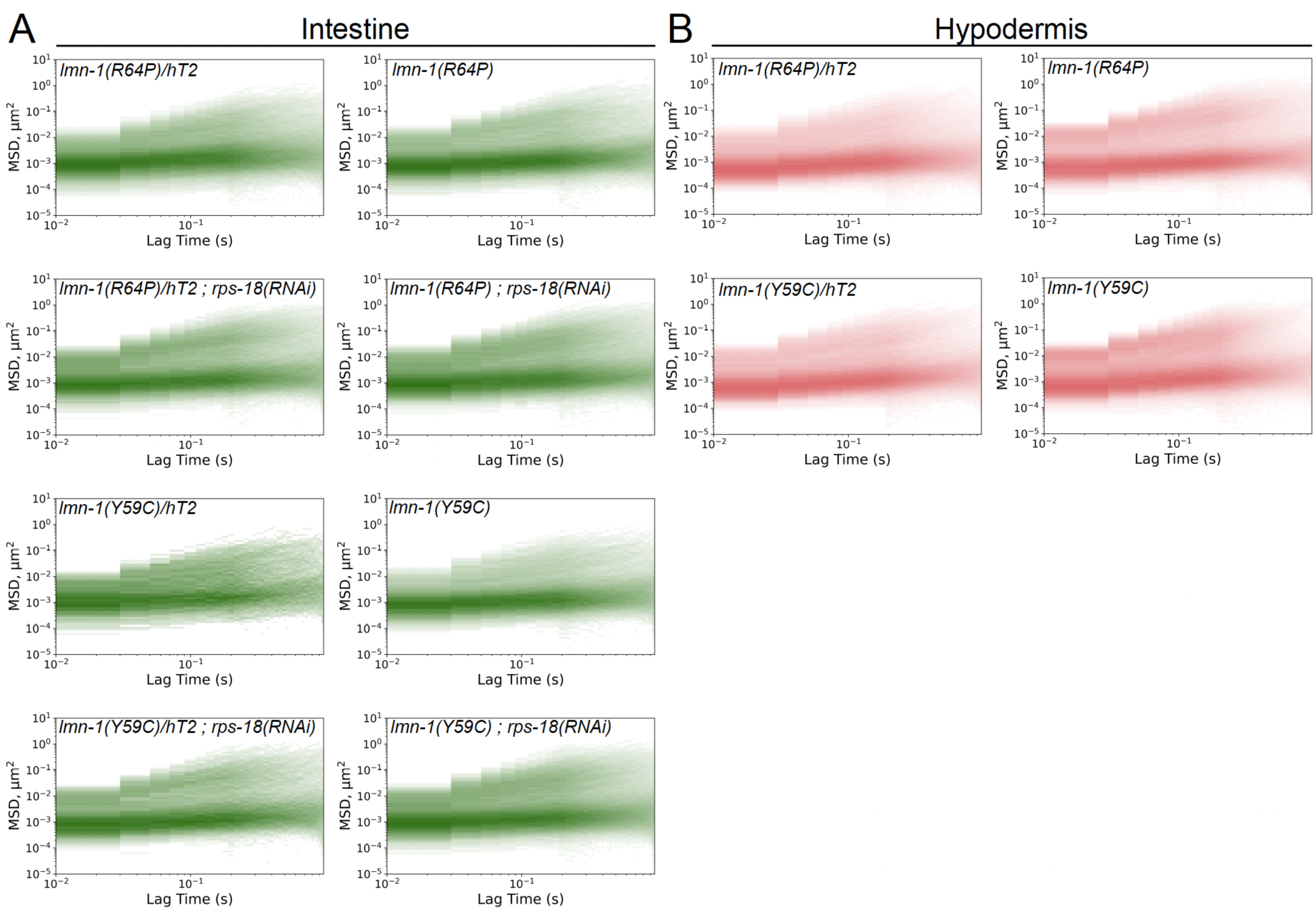
Mean squared displacement (MSD) analysis of each GEM mobility shows minimal population shifts in *lmn-1* mutants. Mean squared displacement (MSD) analysis of GEM mobility over time displayed on a log-log scale for intestinal (A) and hypodermal (B) tissues. Data shown for *lmn-1(R64P)/hT2* heterozygotes, *lmn-1(R64P)* homozygotes, *lmn-1(Y59C)/hT2* heterozygotes, *lmn-1(Y59C)* homozygotes, and corresponding strains treated with *rps-18(RNAi)*. Each plot displays MSD (μm²) versus lag time (s) with color intensity representing track density. Unlike the population shifts observed with *anc-1(e1873)*^10^, these *lmn-1* mutants show relatively subtle changes in the distribution between constrained and unconstrained GEM populations. The overall bimodal pattern is preserved across all conditions, with dense clustering at lower MSD values (constrained population) and more dispersed patterns at higher MSD values (unconstrained population).

**Figure S3.**
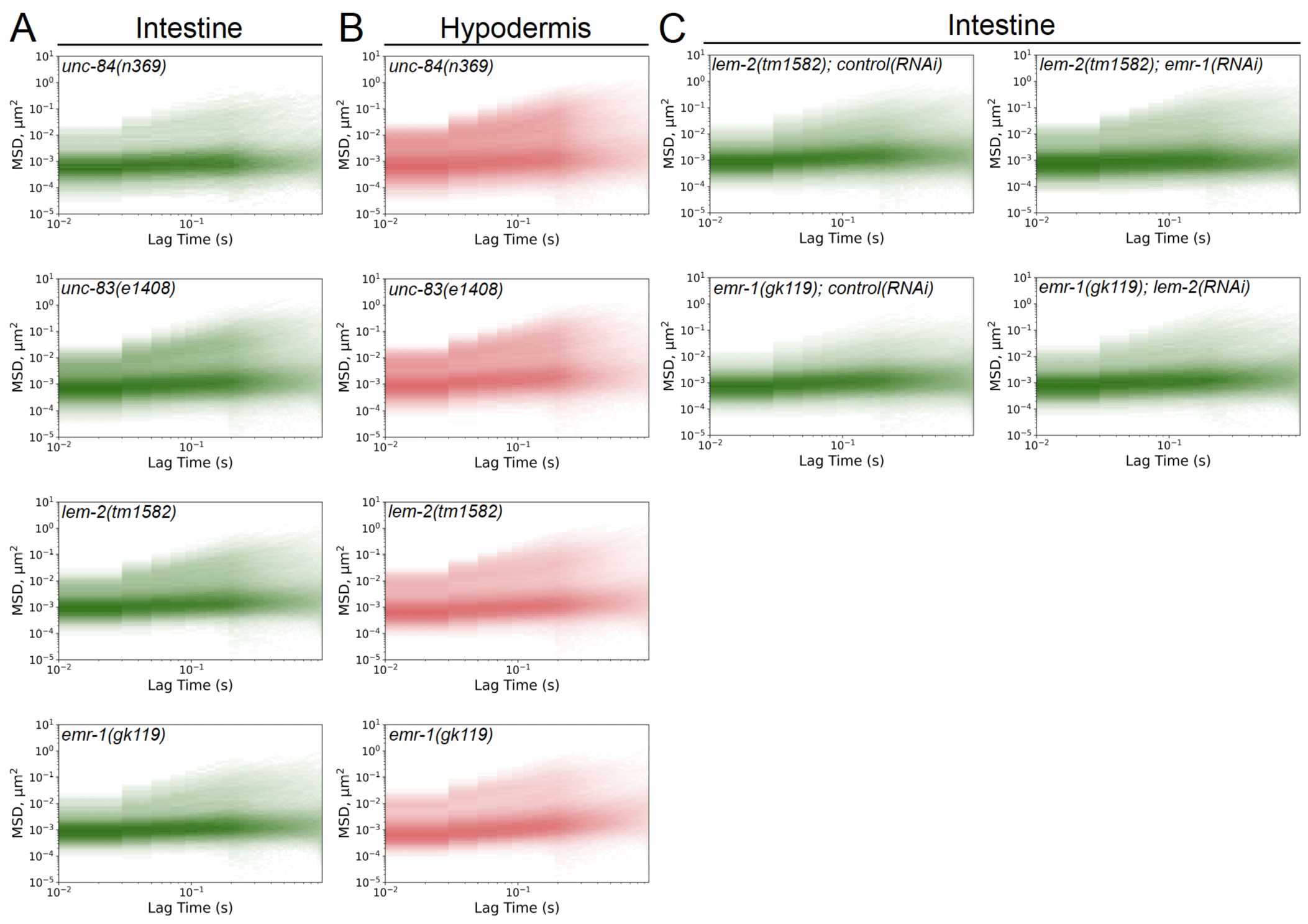
Mean squared displacement (MSD) analysis of each GEM mobility shows minimal population shifts in nuclear envelope protein mutants. Mean squared displacement (MSD) analysis of GEM mobility over time displayed on a log-log scale for intestinal (A, C) and hypodermal (B) tissues. A-B) Data shown for *unc-84(n369)*, *unc-83(e1408)*, *lem-2(tm1582)*, and *emr-1(gk119)* single mutants. C) Data shown for *lem-2(tm1582); control(RNAi), lem-2(tm1582); emr-1(RNAi), emr-1(gk119); control(RNAi),* and *emr-1(gk119); lem-2(RNAi)* conditions. Each plot displays MSD (μm²) versus lag time (s) with color intensity representing track density. The overall bimodal pattern is preserved across all conditions, with dense clustering at lower MSD values (constrained population) and more dispersed patterns at higher MSD values (unconstrained population).

**Movie S1.** Representative movie of a *C. elegans* day 1 young adult carrying the transgene *cSi4[pSL848; pges-1::40nmGEM::eGFP::unc-54 3’UTR] IV*, showing GEMs movement in the intestine. GEMs are pseudocolored in green. *lmn-1(R64P)/hT2* (left) and *lmn-1(R64P)* (right) mutant worms are shown side-by-side. Time bar and scale bar are indicated.

**Movie S2.** Representative movie of a *C. elegans* day 1 young adult carrying the transgene *cSi5[pSL850; psemo-1::40nmGEM::eGFP::unc-54 3’UTR] IV*, showing GEMs movement in the hypodermis. GEMs are pseudocolored in green. *lmn-1(R64P)/hT2* (left) and *lmn-1(R64P)* (right) mutant worms are shown side-by-side. Time bar and scale bar are indicated.

## Author contributions

Conceptualization: XD, DAS, GWGL Investigation: XD, SK, EFG

Methodology: XD, EFG Visualization: XD

Formal analysis: XD, EFG Data curation: XD

Writing & reviewing manuscript: XD, DAS, GWGL

Supervision: DAS, GWGL

Project administration: DAS, GWGL

## Acknowledgements

We thank members of the Starr-Luxton lab for helpful discussions, Dr. Thomas Wilkop and the MCB Light Imaging Facility, Wormbase, and the Center for *C. elegans* Genetics.

